# Functional Screening of Kv7.5 Channel Variants Associated with Neuronal Dysfunction

**DOI:** 10.64898/2026.11.02.763732

**Authors:** Diana V. Hunter, Shefali Vishwakarma, Naseem Givzad, Joshua Liu, Mercedes Sanei, Braden Kersey, Ramon Klein Geltink, Damon T. Poburko, Maksymilian Prondzynski, Tom W. Claydon

## Abstract

The *KCNQ5* gene encodes Kv7.5 channels, which contribute to the regulation of neuronal excitability. Kv7.5 genetic variants associated with neurological dysfunction have been reported, but functional characterization of Kv7.5 variants is limited and causal links to pathogenicity are lacking. We have conducted the first automated patch clamp functional screen of a panel of heterozygous Kv7.5/Kv7.3 channel variants. We tested nine mutations located in diverse locations throughout the Kv7.5 channel and identified some variants with gain-of-function effects (L341I^+/-^, G347S^+/-^, P369R^+/-^, R497C^+/-^), some with loss-of-function effects (V145G^+/-^, W303C^+/-^), and others with no apparent effect (A131V^+/-^, R359H^+/-^, R491H^+/-^). Variants produced a gain of function by left-shifting the voltage-dependence of channel activation alone or in combination with increasing current density. Loss of function occurred as a result of reduced current density. Using these findings, we initiated the first *z*-score classification to guide understanding of Kv7.5 variant pathogenicity, which can be refined as more variants are functionally characterized and new variants are discovered. Lastly, we observed higher sensitivity to channel block by XE991 in L341I^+/-^, G347S^+/-^, P369R^+/-^ gain-of-function variants indicating potential for targeted pharmacogenetic approaches to restore normal Kv7.5 channel function.

## Introduction

Voltage-gated potassium (Kv) channels serve as critical regulators of cellular excitability by conducting repolarizing current in response to changes in the membrane potential. The Kv7 sub-family, Kv7.1-Kv7.5 encoded by the *KCNQ1-5* genes, drives repolarization in neurons, cardiac muscle and vascular smooth muscle (1,2). Kv7.2, Kv7.3, and Kv7.5 are broadly expressed in neurons within the brain and contribute to the M-current, which is a low-voltage activated, non-inactivating, repolarizing current responsible for modulating cell excitability and critical for neuronal electrical and signaling properties (3,4). Kv7.2 and Kv7.3 channels are key components of M-currents in multiple brain regions (5–7), and mutations in the genes encoding Kv7.2 and Kv7.3 channels are linked with neonatal epilepsy (8–10). While Kv7.5 channels are less well studied, they form functional heterotetramers with Kv7.3 subunits and influence neuron excitability (11–15). Indeed, Kv7.5 channels are associated with channelopathies including intellectual disability and epileptic encephalopathy (1,2).

Advances in the understanding of the role of Kv7.5 channels, and proliferation of genetic screening of patients with neurological disorders has unveiled novel Kv7.5 channel variants (16–19). However, causal association with variant pathogenicity is often lacking, which limits the impact on clinical decision making. Moreover, increased accessibility to genetic screening will undoubtedly uncover new Kv7.5 variants of uncertain significance, which presents a hurdle in assigning pathogenicity and patient risk. This emphasizes the need to understand the association between Kv7.5 variant function and pathophysiology, and to create indices for the assignment of pathogenic risk in patients.

The recent development of high content automated patch clamp assays for ion channel variant functional screening has shown significant promise for assigning pathogenicity. For example, high-throughput phenotyping of Kv11.1 (hERG) and Nav1.5 channel variant effects on cell surface expression and channel biophysical gating properties have been used to distinguish pathogenic from benign variants and classify loss-of-function and gain-of-function variant mechanisms (17,20–24). Here, we establish an automated patch clamp screen to examine the impact of Kv7.5 subunit variants on current density and channel function. Investigating a panel of candidate Kv7.5 variants associated with neurological dysfunction, we discovered a range of effects on channel expression and biophysical gating that result in either loss of function or gain of function. This study establishes the first automated patch clamp pathogenic screen of Kv7.5 variants, which promises to improve risk stratification and patient management.

## Materials and methods

### Variant cell line construction

Kv7.5 channel variants were investigated using the reference *KCNQ5* cDNA transcript, NM_001160133.2, and protein sequence, NP_001153605.1. Based on the approach used by Ng et al., (2021)(25), a bicistronic *KCNQ5* plasmid construct was incorporated into the Flp-In T-REx 293 system (Invitrogen, Cat# 978007) to generate variant cell lines. Two copies of the cDNA encoding *KCNQ5*, separated by a linker sequence containing an internal ribosome entry site (IRES), were inserted in the pcDNA5/FRT/TO vector (Invitrogen, Cat# V652020). To enhance translation efficiency, the IRES was placed immediately adjacent to, and directly upstream of, the downstream gene (26). Silent single-base substitutions were introduced into the upstream *KCNQ5* open reading frame to divide the gene into 500-700 bp cassettes, each flanked by unique restriction enzyme sites to facilitate the generation of variants through restriction enzyme-based subcloning. A previously validated hemagglutinin (HA) tag epitope (19,20,27) was inserted into the S1/S2 linker of the upstream *KCNQ5* gene sequence. A C-terminal FLAG tag, analogous to that used previously in K_V_7.2 and K_V_7.3 channels (25,28,29), was appended to the downstream *KCNQ5* gene. The bicistronic wild-type (WT) construct and variant cassettes were synthesized by GeneArt (Thermo Fisher).

WT and heterozygous variant *KCNQ5* Flp-In T-REx 293 cell lines were generated by co-transfecting pcDNA5/FRT/TO containing the desired construct along with pOG44 (Invitrogen, Cat# V600520) to enable Flp recombinase expression and facilitate insertion of the construct into the FRT site within the genome as per the manufacturer’s instructions. Briefly, Flp-In T-REx 293 cells were cultured in DMEM GlutaMAX (Gibco, Cat# 10566016), supplemented with 10% FBS (Life Technologies, Cat# 12483020), 10 µg/mL Blasticidin S HCl (Gibco, Cat# R21001), and 100 µg/mL Zeocin (Gibco, Cat# R25005) at 37°C, 5% CO_2_, and 95% humidity. Once at 40% confluency, cells were co-transfected with 60 ng of pcDNA5/FRT/TO expression plasmid containing the gene construct of interest and 520 ng of pOG44 recombinase plasmid, using Lipofectamine 3000 transfection reagent (Invitrogen, Cat# 3000008) in Opti-MEM (Gibco, Cat# 11058021). Cells were incubated with the transfection reagent in Opti-MEM at 37°C for 5 h, following which the medium was replaced with DMEM GlutaMAX supplemented with 10% FBS and 10 μg/mL Blasticidin S HCl. After 24 h, the medium was exchanged for DMEM GlutaMAX containing 10% FBS, 10 μg/mL Blasticidin S HCl, and 100 μg/mL Hygromycin B (Gibco, Cat# 10687010), which was replaced every 4 days to eliminate cells sensitive to the antibiotic selection pressure. Upon formation of antibiotic-resistant WT or variant *KCNQ5* expressing cell colonies, cells lines were expanded by lifting with TrypLE Express Enzyme (no phenol, Gibco, Cat# 12604021) and re-seeding. To induce *KCNQ5* gene expression, cell lines were grown to 40% confluency before addition of 1 µg/mL tetracycline (Gibco, Cat# A39246) for 48 h prior to experiments.

### Transient transfection with KCNQ3

To facilitate trafficking of Kv7.5 channels in HEK cells (11), Flp-In T-Rex 293 cells stably expressing WT or heterozygous variant *KCNQ5* were transiently transfected with WT *KCNQ3* (a kind gift from Dr. Harley Kurata) and GFP 48 h ahead of characterization using the high efficiency ExPERT ATx electroporator (MaxCyte). Electroporation was performed according to the manufacturer’s directions using either OC-25×3 or R-50×8 process assemblies. Briefly, WT or heterozygous variant *KCNQ5* Flp-In T-Rex 293 cells were detached using TrypLE Express Enzyme, washed and resuspended in MaxCyte Electroporation Buffer (Cytiva, Catalog # EPB-1) at 1 × 10^8^ cells/ml, and mixed with 50 ng/μL pEGFP-N1 and 150 ng/μL *KCNQ3* in pcDNA3.1(-). Loaded assemblies were then electroporated using the manufacturer pre-defined parameters for HEK cell transfection. Following transfection, cells were incubated with DNase I in DMEM for 30 min at 37°C, and then incubated for 48 h in DMEM GlutaMAX with 10% FBS and 1 µg/mL tetracycline to allow for both *KCNQ3* and *KCNQ5* protein expression.

### Electrophysiological recording

Whole-cell patch-clamp of WT or heterozygous variant Kv7.5/Kv7.3 channels was conducted using NPC-384T S-type chips and the SyncroPatch 384-well automated planar patch clamp platform (Nanion Technologies, Cat# 22 1101) under voltage-clamp conditions. Cells were dissociated using TrypLE Express Enzyme and resuspended in CHO-S-SFM-II media (Gibco, Cat# 31033020C). The cell suspension was triturated and centrifuged at 150 g for 8 min after which the cell pellet was resuspended in external recording solution (External Standard; composition in mM: 140 NaCl, 4 KCl, 2 CaCl_2_, 1 MgCl_2_, 5 Glucose, 10 HEPES to pH 7.4, Nanion Technologies Cat# 08 3001) at a final concentration of 1 - 2 × 10^5^ cells/ml. Cells were loaded onto the SyncroPatch 384 cell hotel, which was incubated at 10°C and rotated at 300 rpm for 1 h before recordings. User-defined voltage clamp protocols were developed using the Biomek i5 and PatchControl 384 Version 3.4.0 platform. Cell catch was performed at-40 mBar in divalent-free external solution (External [-] Ca^2+^ [-] Mg^2+^; composition in mM: 140 NaCl, 4 KCl, 5 glucose, 10 HEPES to pH, Nanion Technologies Cat# 08 3003) and KF110 internal solution (Internal KF110; composition in mM: 110 KF, 10 KCl, 10 NaCl, 10 EGTA, 10 HEPES to pH 7.2, Nanion Technologies Cat# 08 3007) or KF40 internal solution (composition in mM: 80 KCl, 40 KF, 5 EGTA, 4 Na-ATP, 2.5 CaCl_2_ to pH 7.2 supplemented with 1 μM diC8-PIP_2_) for 2 min followed by external solution exchange with seal enhancer solution (External Standard Ca 10; composition in mM: 130 NaCl, 4 KCl, 10 CaCl_2_, 1 MgCl_2_, 5 Glucose, 10 HEPES to pH 7.4, Nanion Technologies Cat# 08 3012). Following two washes with external recording solution the whole cell configuration was achieved by applying-100 mBar pressure for 2 min. To measure channel activation properties, currents were elicited from a holding potential of-80 mV by applying a series of 1.5 s voltage steps ranging from-110 mV to +40 mV in +10 mV increments followed by a 500 ms step to 0 mV for tail current analysis (step frequency, 0.3125 Hz). Following control recordings, the K_V_7 channel blocker, XE991 (Sigma, Cat# X2254) (30), was applied. Following a 20 s voltage step to 0 mV to enable access to its binding site (31), channel activation was measured again using the above voltage step protocol. Recordings were made from each well following application of 1 μM XE991 and 30 μM XE991. Experiments were conducted at 25°C and series resistance compensation was set to 80%.

### Flow Cytometry

Cells were fixed using 4% paraformaldehyde (PFA) for 10 min at room temperature, washed in phosphate buffered saline (PBS, VWR, Cat# CA16777-251), and stored at 4 °C for flow cytometry. Fixed cells were blocked using 2% FBS in PBS with 2 mM EDTA for 1 h at room temperature and then stained with 1:1000 anti-HA antibody primary conjugated to Alexa 647 (Invitrogen, Cat# 740005MP647) for 1 h at room temperature. A sample of WT *KCNQ5-HA-tag* expressing cells without *KCNQ3* transfection were compared to a sample of the WT KCNQ5-HA-tag bicistronic Flp-In T-Rex 293 cells after MaxCyte electroporation with pCDNA3.1(-) KCNQ3 and pEGFP-N1. Samples were acquired on a BD LSRFortessa X-20 Cell Analyzer, with a at least 0.5 x 10^5^ events collected for the debris-and doublet-excluded population. Debris was excluded based on forward scatter area (FSC-A) and side scatter area (SSC-A). Doublets were discriminated based on forward scatter area and height (FSC-A vs. FSC-H), and side scatter area and height (SSC-A vs. SSC-H). Events were subsequently gated on the fluorophore-positive population using fluorescence-minus-one and unstained control. Data were analyzed using FlowJo v10 (BD Biosciences).

### Data analysis

Electrophysiological recordings were analysed using DataControl 384 Version 3.2.1 (Nanion Technologies). A series of quality control parameters were used to create inclusion/exclusion criteria across wells. All quality control parameters were required to be met for a well to be included. Inclusion criteria required wells to maintain a seal resistance >400 MΩ throughout all recordings and drug applications, a cell capacitance of 1-25 pF. Analyses were conducted on data without leak subtraction.

Current density was calculated from the isochronal peak current amplitude at the end of the voltage step to +40 mV normalized to the membrane capacitance in all wells that met the inclusion criteria. Conductance-voltage relationships were generated from wells with >200 pA peak tail current amplitude that was >85% inhibited by application of 30 μM XE991. Peak tail currents following each voltage step (*G*) were normalized to the peak tail current following the +40 mV step (*G_max_*) and were fitted using a Boltzmann function:

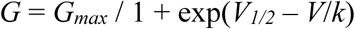

where V is the test potential, *V_1/2_* is the half-maximal activation potential, and *k* is the slope factor. The time course of channel activation was measured by fitting the rising phase of current in response to a voltage step to 0 mV to a single exponential function:

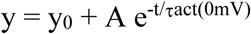

where y is the normalized response, A is the amplitude, and τ_act(0mV)_ is the time course. A *z*-score-based classification of variant effects based on that described by Ng et al. (2021) was used (25). To calculate *z*-scores for variant effects on current density, data were first square root transformed to help approximate a normal distribution and the variant current density was expressed relative to the distribution of the WT data using:

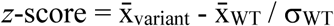

where x̄_variant_ is the mean current density in the given variant, x̄_WT_ is the mean current density of WT, and σ_WT_ is the standard deviation of the WT current density. A similar approach was used to calculate *z*-scores for the effect of each variant on the *V_1/2_* of activation and the activation tau. In these cases, the data were normally distributed and did not require root transformation.

### Statistics and data representation

Statistical testing and data representation were performed using GraphPad Prism 10.4.2. Data were tested for normality using the Shapiro–Wilk normality test, and comparisons between measures from WT and Kv7.5 variants were made using a one-way ANOVA with a Dunnett’s or Kruskal-Wallis with a Dunn’s post hoc test for parametric and nonparametric data, respectively. A *P*-value of <0.05 was used to indicate a significant difference. Box plots indicate the interquartile range, where the box limits represent the 25% and 75% percentiles; the middle line reflects the median; and the whiskers represent the 5% and 95% percentiles of the data. Symbols superimposed on box plots represent data from individual cells. In other plots, symbols reflect the mean ± standard error of the mean. In plots with grey shading, the darker shading represents the SEM of WT and the lighter shading indicates the 95% confidence intervals. Variants with mean values outside of the 95% confidence interval of WT were considered to be significantly different from WT. Values for *n* reflect the sample size.

## Results

### Development of an automated patch clamp screen for Kv7.5 channels

Initial automated patch clamp electrophysiological recordings of channel function from inducible stably transfected WT bicistronic Kv7.5 Flp-In T-REx 293 cells at 37°C revealed currents with an unanticipated voltage-and time-dependence (Figure 1A). Previous observations suggest that rundown of channel function may be a significant contributing factor to the current phenotype observed in these initial recordings (32). Consequently, optimization of recordings conditions included alteration of the internal recording solution to include 40 mM KF (lowered from 110 mM), 1 μM diC8-PIP_2_, and 2.5 mM CaCl_2_, recording currents at 25°C, and co-expressing Kv7.3 channel subunits. These optimized conditions produced robust and reproducible Kv7.5 currents that eliminated rundown within voltage steps and throughout the duration of the experiment (Figure 1B). Immunodetection of membrane surface Kv7.5 channel protein using an S1-S2 extracellular linker HA epitope revealed that co-expression of Kv7.3 was particularly important for surface expression of Kv7.5 (Figure 1C) as has been reported previously (20). Fits of the voltage-dependence of activation of Kv7.5/Kv7.3 heteromultimer channels recorded with these optimized conditions with a Boltzmann function yielded *V_1/2_* and *k* values of-27.4 ± 3.7 mV and 17.4 ± 0.65 mV, respectively, while the mean time constant of activation at 0 mV (τ_act(0mV)_) was 97.3 ± 7.1 ms (Figure 1D). These biophysical properties indicate that, measured under these conditions, Kv7.5 channel function in our automated patch clamp assay is similar to that reported previously in other heterologous cell systems (16–19).

**Figure 1.**
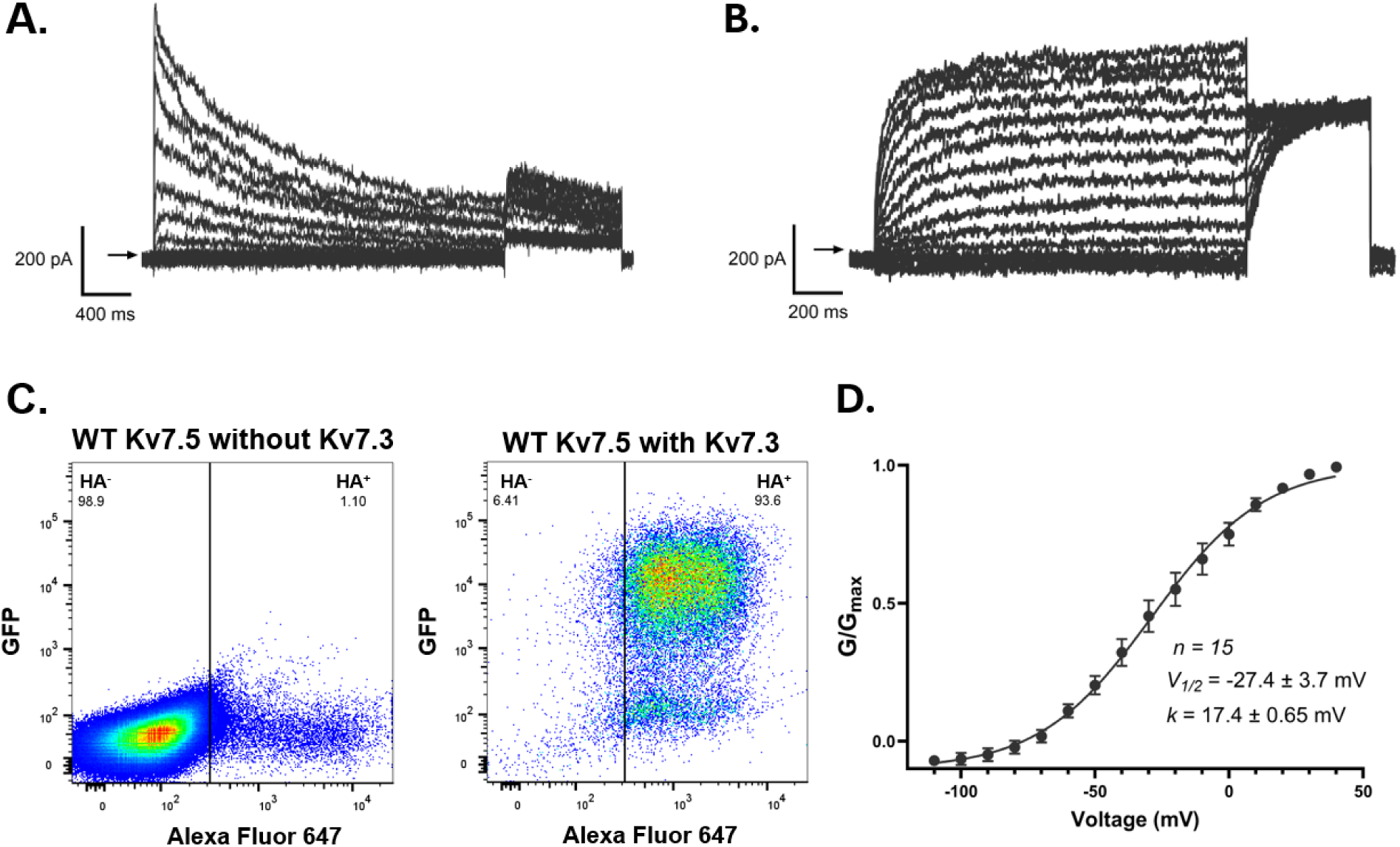
Optimization of automated patch clamp conditions for Kv7.5 recordings. **(A)** Representative current family recorded in response to the voltage protocol shown from WT Kv7.5 channels at 37°C. **(B)** Representative current family recorded from WT Kv7.5 channels with co-expression of Kv7.3 and measured in optimized recording conditions. **(C)** Scatter plots from flow cytometry showing the proportion of cells staining positive (HA+) and negative (HA-) for HA-tagged Kv7.5 in the absence (*left*) or presence (*right*) of Kv7.3+GFP co-expression. **(D)** Mean conductance-voltage relationship from WT Kv7.5/Kv7.3 channels fitted with a Boltzmann function. Mean values from fits to the WT data are shown. Data are presented as mean ± SEM.

### Functional screening of a panel of Kv7.5 channel variants

Having optimized conditions to record WT Kv7.5 currents using automated patch clamp, we investigated the functional impacts of a panel of heterozygous Kv7.5 variants. Flp-In T-REx 293 cell lines expressing the heterozygous variants, A131V^+/-^, V145G^+/-^, W303C^+/-^, G347S^+/-^, R359H^+/-^, P369R^+/-^, R491H^+/-^, and R497C^+/-^ were generated and studied, each with transient transfection of Kv7.3. This panel contains a range of Kv7.5 variants that are reported as likely pathogenic, or variants of uncertain significance (33,34). The variants are distributed across multiple structural and functional regions of the Kv7.5 channel, including the voltage-sensing domain, pore region, Helix A, and intrinsically disordered regions of the C-terminus (Figure 2A). The 384-well automated patch clamp chip layout was designed to incorporate electrophysiological characterization of the function, biophysical gating properties, and sensitivity to the Kv7 channel blocker, XE991, of WT and each variant across ≥32 individual wells each. All wells were filtered using quality-control criteria based on seal resistance (≥400 MΩ), whole cell capacitance (1-25 pF), and recording stability throughout the duration of the experiment, and the number of wells passing quality control parameters was relatively consistent (49.3 ± 2.26 %) across the panel.

**Figure 2.**
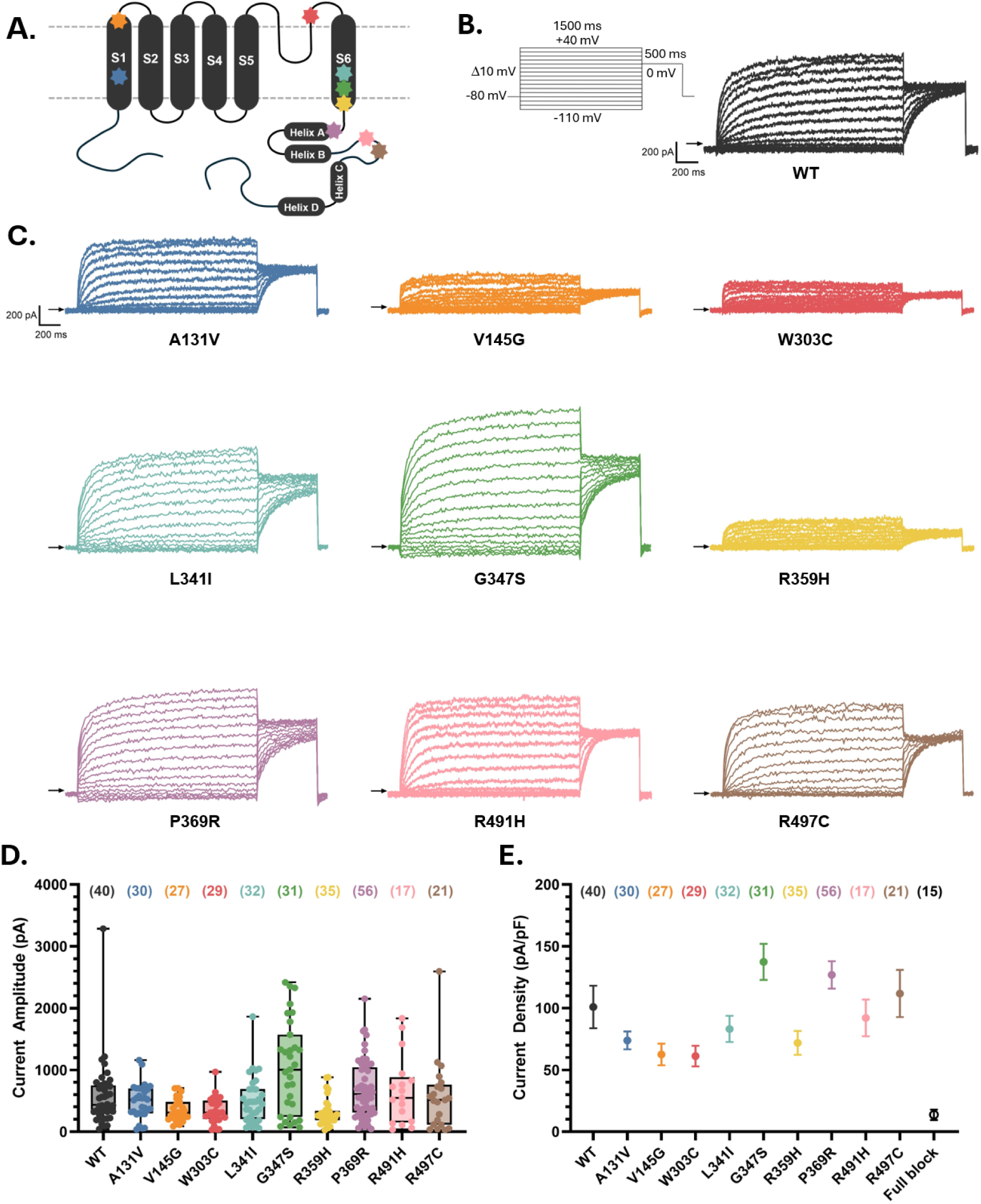
Characterization of Kv7.5/Kv7.3 channel function and surface expression. **(A)** Schematic of Kv7.5 channel topology showing location of each variant studied. **(B)** Representative whole-cell current trace family recorded from WT Kv7.5 cells co-expressing Kv7.3 at 25°C in response to the voltage protocol shown. **(C)** Representative current trace family recorded from each Kv7.5 variant studied. **(D)** Peak current amplitudes recorded from WT and variant Kv7.5/Kv7.3 channels in all cells that passed quality control measures. Numbers in parenthesis indicate sample size. **(E)** Plot of the mean ± SEM current density in WT and variant Kv7.5/Kv7.3 channels. The mean ± SEM current density in WT following full block with 30 μM XE991 is also plotted to indicate background current levels.

A representative example of currents recorded from WT and each heterozygous Kv7.5 variant in the panel is shown in Figure 2B and C. The peak isochronal current amplitude measured at the end of the voltage step to +40 mV from all WT and Kv7.5 variant cells that passed quality control measures is shown in Figure 2D. The mean peak current amplitude was 584 ± 84 pA in WT cells (n = 40), and this measure appeared to be reduced in some variants and increased in others. To account for individual cell capacitance and to enable quantification of the relative function of each channel type, the mean current density was calculated and is plotted in Figure 2E. Statistical comparison across the panel indicated that G347S^+/-^ (136 % of WT function) and P369R^+/-^ (126 % of WT function) increased the current density compared with WT (Kruskal-Wallis non-parametric test, p = 0.0142 for G347S^+/-^ and p = 0.0268 for P369R^+/-^). Although they did not reach statistical significance, the current density in V145G^+/-^ and W303C^+/-^ was 62 and 61 % that of WT, respectively. The current density in WT channels following full channel block by 30 μM XE991 indicates background expression levels (13.7 ± 4.2 pA/pF) in our experiments. These findings indicate that two Kv7.5 variants have gain-of-function effects in the heterozygous setting.

### *Incorporating a* z*-score categorization of Kv7.5 variant function*

To provide a standardized quantitative measure of the impact of Kv7.5 variant activity across the panel, square root transformed current density values were normalized to WT values to generate a *z*-score-based activity classification as done previously (35). In this approach, WT Kv7.5 was assigned a reference value of 0, with each variant *z*-score used to determine the degree of functional perturbation. Figure 3A ranks the current density *z*-scores in each heterozygous variant and highlights the 95% confidence intervals of the WT data (light grey;-0.32 – 0.32). These data indicate a spread of functional effects with G347S^+/-^ (*z*-score = 0.49 ± 0.14) and P369R^+/-^ (*z*-score = 0.38 ± 0.10) introducing a significant gain-of-function effect in the heterozygous setting. Again, loss-of-function heterozygous variant effects, e.g. V145G^+/-^ (*z*-score =-0.33 ± 0.11) and W303C^+/-^ (*z*-score =-0.35 ± 0.11), were more difficult to discern. We reasoned that the sensitivity to detect loss of function may be impacted because our automated patch clamp approach requires transient transfection of Kv7.3 to facilitate surface expression of Kv7.5 subunits. Although transfection efficiencies with Kv7.3 were high (81.1 ± 4.2 %), some cells may lack Kv7.3, which would reduce Kv7.5 current density. To account for this, we recalculated the current density *z*-scores using only cells with >200 pA peak current amplitude (Figure 3B). With this cut-off applied, variant *z*-scores ranged from-0.64 to 0.57, with a significantly different from WT (*z*-score 95% confidence intervals =-0.35 – 0.35) gain of function produced by G347S^+/-^ (*z*-score = 0.57 ± 0.13), R497C^+/-^ (*z*-score = 0.40 ± 0.20), and P369R^+/-^ (*z*-score = 0.39 ± 0.10), and significant loss of function produced by W303C^+/-^ (*z*-score =-0.42 ± 0.13) and V145G^+/-^ (*z* score =-0.64 ± 0.089).

**Figure 3.**
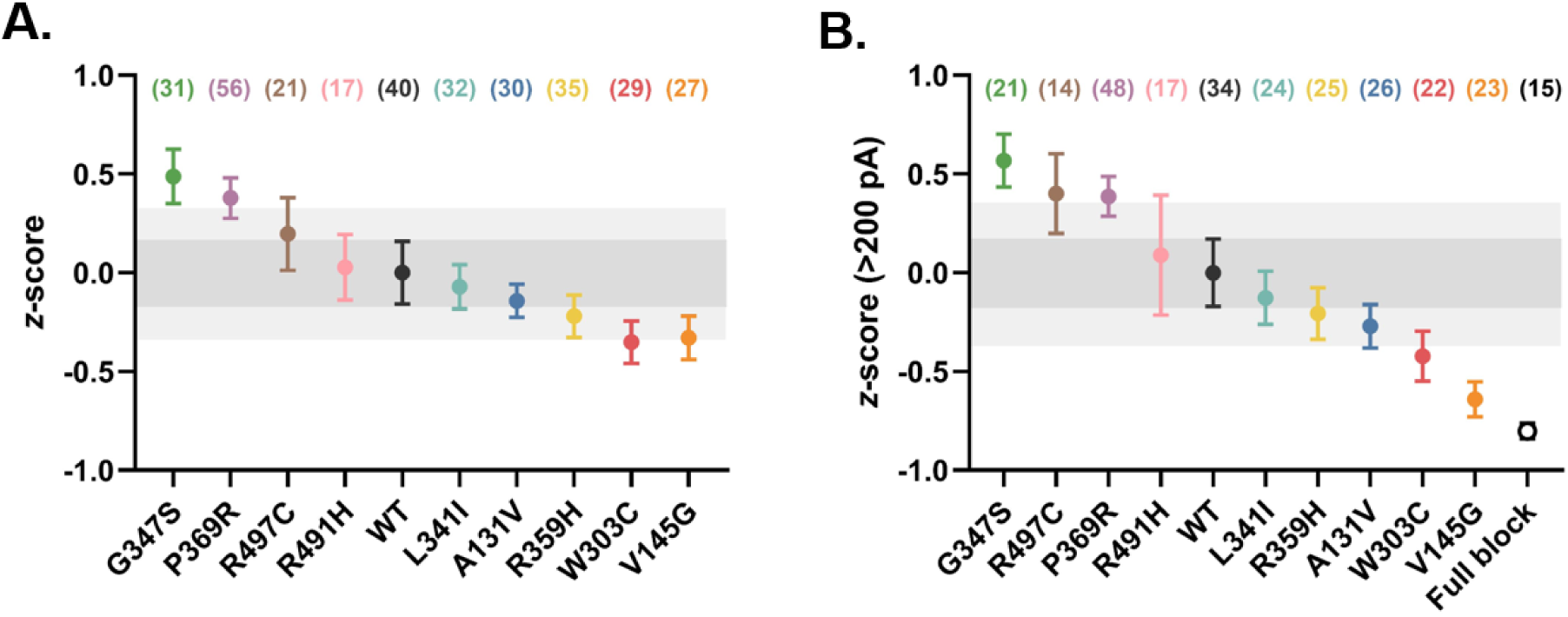
*z*-score representations of Kv7.5 variant effects on current density. **(A)** *z*-scores calculated for Kv7.5 variant cells that passed quality control, in order from largest to smallest score. **(B)** *z*-scores calculated for Kv7.5 variant cells with ≥200 pA peak current amplitude. Data are presented as mean ± SEM. WT is set to zero with dark grey shading showing ± SEM range and light grey showing ± 95% confidence interval for WT. Numbers in parenthesis indicate sample size.

### Kv7.5 variant effects on biophysical properties

In addition to effects on current density, Kv7.5 channel variants may influence neuronal function by altering the biophysical gating properties of the channel. Figure 4 shows the effects of each heterozygous variant on the voltage-dependence of channel activation and the time course of channel opening. Normalized tail currents were used to plot mean conductance-voltage relationships for each variant, which were fitted with a Boltzmann function (Figure 4A). Mean values for *V_1/2_* and *k* derived from Boltzmann fits of WT and variant channels are shown alongside the fits in Figure 4A and the *V_1/2_* values measured from individual cells are plotted in Figure 4B. Time constant values derived from single exponential fits of current activation at 0 mV in each variant (τ_act(0mV)_) are shown in Figure 4C. In a similar approach to quantifying perturbations in current density, we calculated *z*-scores describing the effects of the panel of variants on the *V_1/2_* of activation and on activation kinetics (Figure 4C and E) to describe variant-induced perturbations in biophysical gating properties. These data indicate that the *V_1/2_* was significantly left-shifted compared with WT (*z*-score 95% confidence interval range =-0.55 - 0.55) in the heterozygous variants, L341I^+/-^ (*z*-score =-0.97 ± 0.28), G347S^+/-^ (*z*-score =-1.07 ± 0.14), and P369R^+/-^ (*z*-score =-1.13 ± 0.28). The τ_act(0mV)_ value was significantly greater compared with WT (95% confidence interval range =-0.55 – 0.55) in the heterozygous variants, G347S^+/-^ (*z*-score = 0.88 ± 0.21), R497C^+/-^ (*z*-score = 0.70 ± 0.41), R359H^+/-^ (*z*-score = 0.67 ± 0.46), and P369R^+/-^ (*z*-score = 0.57 ± 0.19). It is notable that the *z*-score values reflecting the change in the *V_1/2_* from WT indicate larger perturbation in function than the effect on current density values in the gain-of-function variants, G347S^+/-^ and P369R^+/-^.

**Figure 4.**
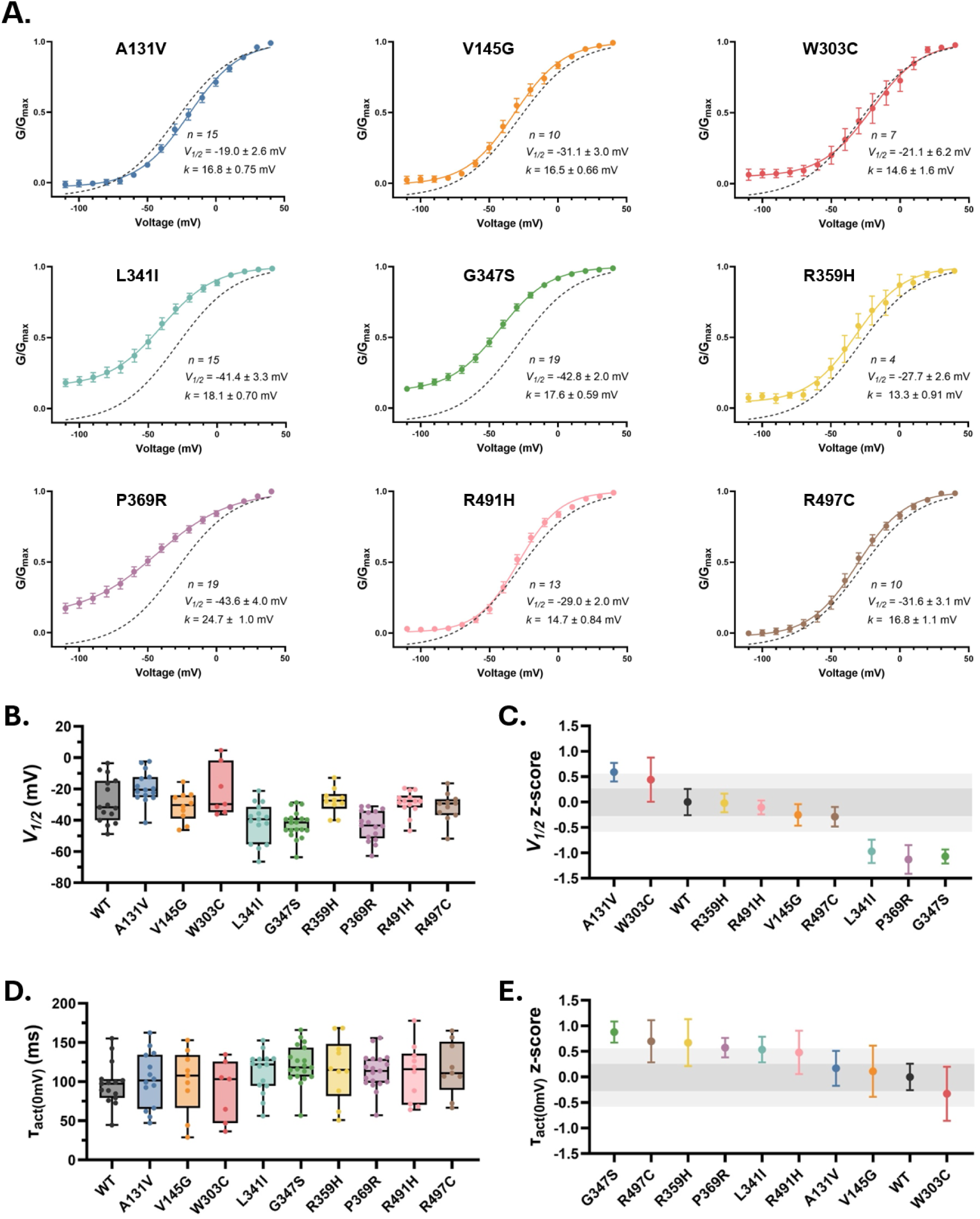
Biophysical characterization of Kv7.5 variants. **(A)** Mean conductance-voltage relationships of variant Kv7.5/Kv7.3 channels. The mean relationship measured in WT from Figure 1D is also plotted as a grey dashed line for reference. **(B)** Plot showing the half-activation voltage (*V_1/2_*) for WT and variant Kv7.5/Kv7.3 channels in all cells. **(C)** *z*-score representations of Kv7.5 variant effects on the *V_1/2_* of activation. Variants are ranked according to effect size. **(D)** Plot of the kinetics of channel activation (*τ*_act(0mV)_) for WT and variant Kv7.5/Kv7.3 channels in all cells. **(E)** *z*-score representations of Kv7.5 variant effects on the *τ*_act(0mV)_. Variants are ranked according to effect size. Data are presented as mean ± SEM. For *z*-score plots, WT is set to zero with dark grey shading showing ± SEM range and light grey showing ± 95% confidence interval for WT.

### Drug sensitivity of the Kv7.5 variant panel

The compound, XE991, is a potent and selective inhibitor of Kv7.2-Kv7.5 channels that preferentially blocks activated channels (31). To investigate the selective inhibition of gain-of-function variants, we measured the extent of block induced by 1 μM XE991 in the heterozygous Kv7.5 variant panel. Following a 20 s step to 0 mV to enable XE991 to reach its binding site, we observed that 1 μM XE991 inhibited peak outward current during the +40 mV voltage step in WT channels by 65.8 ± 4.3 % (Figure 5A and B). XE991 inhibited channel current without shifting the voltage-dependence of channel activation (Figure 5C), indicative of an open channel block mechanism as reported previously (31). Figure 5D compares the extent of inhibition of the panel of heterozygous Kv7.5 variants and shows that some variants affected the sensitivity of channel block by 1 μM XE991. The extent of block was statistically reduced by the V145G^+/-^ (p = 0.0107), L341I^+/-^ (p = 0.0108) and P369R^+/-^ (p = 0.0025) variants (Kruskal-Wallis with Dunn’s post hoc test). As observed in WT channels, the application of XE991 did not alter the *V_1/2_* of channel activation in any of the panel variants.

**Figure 5.**
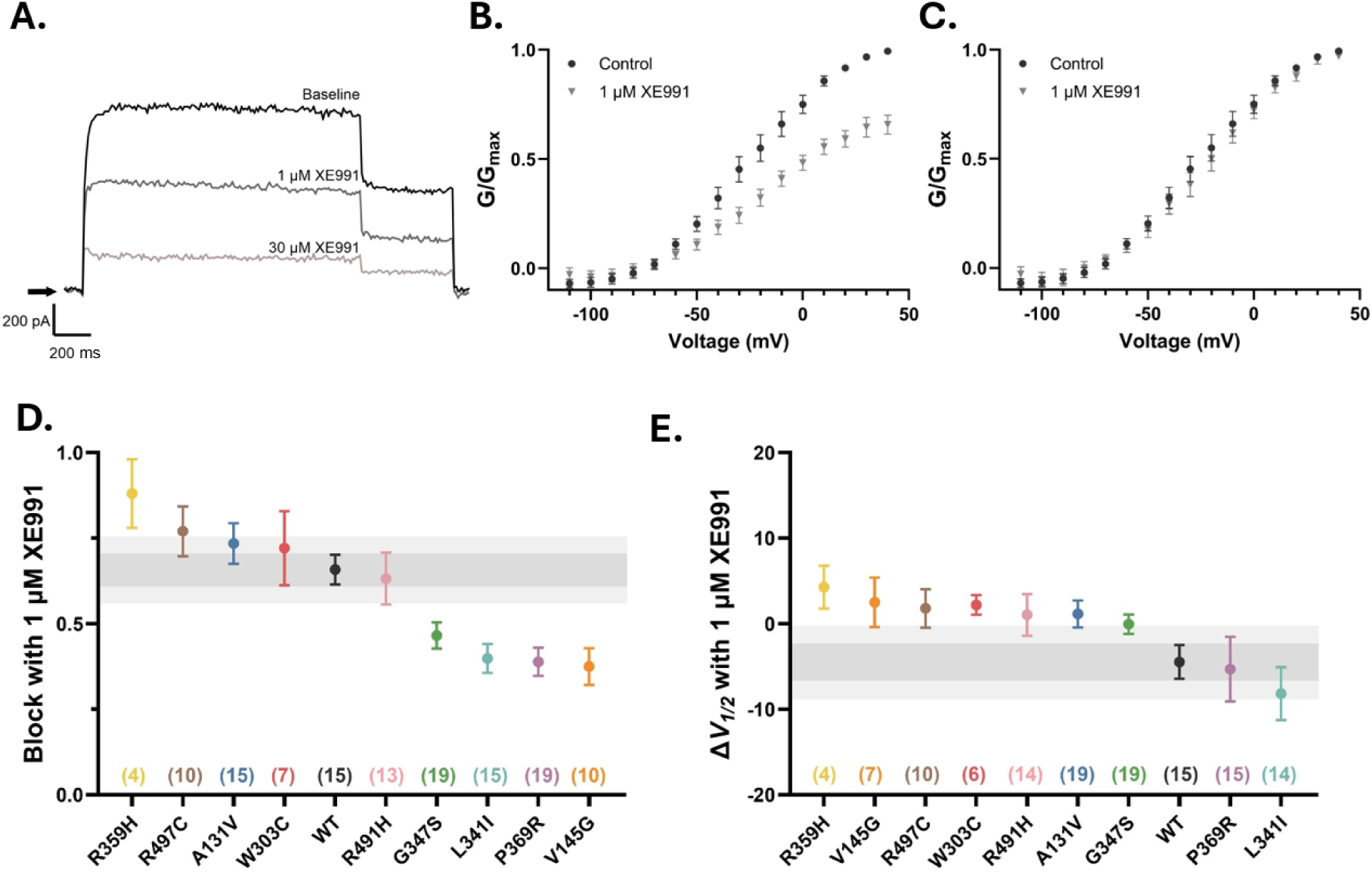
Effect of XE991 on Kv7.5/Kv7.3 variant channel function. **(A)** Representative Kv7.5 WT channel whole-cell current traces recorded at +40 mV from the same cell during control conditions and following application of 1 µM and 30 µM XE991. **(B)** Mean WT Kv7.5/Kv7.3 maximal conductance during control and with 1 µM XE991 normalized to that during control to highlight extent of block. **(C)** Mean WT Kv7.5/Kv7.3 maximal conductance during control and with 1 µM XE991 normalized to their own condition to highlight XE991-induced shifts in the voltage dependence of activation. **(D)** Fractional maximal conductance following 1 µM XE991 application highlight Kv7.5 variant effects on channel sensitivity to XE991 block. Numbers in parenthesis indicate sample size. **(E)** Mean change in the *V_1/2_* of activation for WT and Kv7.5 variants caused by the application of 1 µM XE991. Data are presented as mean ± SEM. For *z*-score plots, WT is set to zero with dark grey shading showing ± SEM range and light grey showing ± 95% confidence interval for WT.

## Discussion

Heterozygous Kv7.5 channel variants are associated with neurological disorders, but a lack of functional characterization of variants, and of causal links to pathogenicity limits patient management. This study presents the first automated patch clamp screen of heterozygous Kv7.5/Kv7.3 channel variant function. We explored the effects of nine mutations located in diverse locations throughout the Kv7.5 channel on Kv7.5/Kv7.3 channel expression and biophysical properties. We found that some variants had gain-of-function effects (L341I^+/-^, G347S^+/-^, P369R^+/-^), while others had loss-of-function effects (V145G^+/-^, W303C^+/-^). Gain of function arose from biophysical effects of variants on channel gating (left-shifted voltage-dependence of channel activation) alone or in combination with channel expression (increased current density). Loss of function occurred as a result of reduced current density. These findings guided the initiation of a *z*-score based index for classification of Kv7.5 variant pathogenicity. We also demonstrated selective inhibition of L341I^+/-^, G347S^+/-^, P369R^+/-^ gain-of-function variants using a Kv7 channel blocker indicating potential for targeted pharmacogenetic approaches to restore normal Kv7.5 channel function.

### Establishing an automated patch clamp screen of Kv7.5 variant function

It is challenging to meet the need for broad Kv7.5 variant functional characterization using conventional manual patch clamp. Automated patch clamp screens, on the other hand, offer significant high content opportunities. However, complexities related to heteromultimeric Kv7 channel subunit assembly and rundown of function due to PIP_2_ depletion present hurdles that must be overcome (32,36). Here, we have established favourable conditions with which to record and screen variant Kv7.5 channel function, which include co-expression with Kv7.3 subunits and modification of the intracellular recording solution to include PIP_2_ and calcium supplementation and reduced fluoride concentration. The most significant factor was co-expression of Kv7.3. While Kv7.5 channel subunits can function as homotetramers, current density is greatly enhanced by the presence of Kv7.3 subunits, which form heterotetramers with Kv7.5 in heterologous expression systems and native tissues (37,38). It is therefore important to investigate the effects of heterozygous Kv7.5 variants in the presence of Kv7.3 to aid translation of their physiological consequences. The need for inclusion of Kv7.3 in Kv7.5 functional screens is highlighted in the present study, which shows transformed robust functional Kv7.5/Kv7.3 currents. Co-assembly of Kv7.5 with Kv7.3 subunits also confers improved resistance to channel rundown due to PIP_2_ depletion (39,40), and along with increased free intracellular calcium, which stabilizes calmodulin-mediated PIP_2_ channel interaction (41), and supplementation with exogenous PIP_2_, this produced reproducible currents with little rundown of function in our automated patch clamp. We also found it critical to reduce intracellular fluoride, which is utilized in automated planar patch clamp systems to enhance gigaohm seal formation (42). Fluoride activates phospholipase-C-dependent hydrolysis of PIP_2_ and inhibits phosphatidylinositol-4-kinase-dependent resynthesis of PIP_2_ (39,43), and contributes to Kv7.5 channel rundown, and we found that lowering the intracellular KF to 40 mM reduced channel rundown without greatly compromising the ability to form tight gigaohm seals. Future studies to reduce/prevent rundown in Kv7 channels may benefit from the use of fluoride-free recording chips that have recently been developed (42).

### Towards the establishment of a z-score based Kv7.5 variant pathogenicity index

Our findings are the first using automated patch clamp to report the effects of a series of heterozygous Kv7.5 variants co-expressed with Kv7.3 and indicate that variants perturb channel function by altering current density and/or gating properties. Current density measurements are often used as a proxy for functional channel expression and in our panel Kv7.5 currents were measured at a strongly depolarized voltage that reflected maximal channel conductance in each variant. We established *z*-scores to describe the effects of each Kv7.5 variant on current density relative to the standard deviation of WT as done previously for variant screens in other channel types (21–23). Our *z*-score findings indicate that G347S^+/-^ (131% increase over WT), R497C^+/-^ (127% increase over WT), and P369R^+/-^ (121% increase over WT) exhibit gain-of-function effects on current density, while V145G^+/-^ (58.5% less than WT) and W303C^+/-^ (59.0% less than WT) produce loss-of-function effects, and A131V^+/-^, L341I^+/-^, and R369H^+/-^ have no effect. These findings are consistent with previous observations in homozygous G347S, P369R, and V145G channels (16–18), which serve to validate our automated patch assay. However, our findings extend these observations to include previously unreported variants and measurement in the heterozygous setting, which is the typical clinical presentation. Furthermore, our *z*-score approach enables a quantitative description of variant effects relative to the observed variance in WT function.

In addition to effects on current density, we observed variant-induced effects on the biophysical gating characteristics of Kv7.5 channels. L341I^+/-^, G347S^+/-^, and P369R^+/-^ produced gain-of-function effects by shifting the voltage-dependence of activation to the left, while G347S^+/-^, R497C^+/-^, and R359H^+/-^ produced a mild slowing of activation. Of note, L341I^+/-^, G347S^+/-^, and P369R^+/-^ channels did not close even at strongly hyperpolarized membrane voltages indicating constitutive activation in these variant channels. Such biophysical gating changes may relate to altered PIP_2_ sensitivity and channel open probability resulting in enhanced current density measurements (18). These alterations would be expected to have profound effects on excitability in neuronal networks, since the M-current is the only sustained current that flows near the action potential threshold impeding repetitive action potential firing and dampening responses to depolarizing inputs (1,44). However, since variants may alter the voltage-dependence and time course of channel activation independently of effects on current density, these robust gain-of-function effects on biophysical characteristics highlight the importance of incorporating gating effects and current density effects into future Kv7.5 variant functional screens.

### The need for Kv7.5 variant z-score refinement

While our findings indicate significant effects in variants whose *z*-score lies outside of the 95% confidence interval of the WT data set, it is striking that the magnitude of *z*-scores observed for our Kv7.5 variants is lower than that reported in other channel types (22,35). In these studies, a threshold *z*-score of ≥-2 to +2, which reflects two standard deviations from control, is often used to distinguish perturbed from normal function. One possible interpretation of this is that the physiological consequences of Kv7.5 channel dysfunction are profound, such that the lower *z*-scores observed here result in greater neuronal functional perturbation compared with the perturbation in function associated with a similar *z*-score in cardiac hERG or *SCN5A* channels. Another possibility is that the reduced impact of Kv7.5 variants on channel expression is related to the co-assembly with Kv7.3, which enhances trafficking of Kv7.5 subunit-containing heterotetramers. In the presence of a Kv7.3-driven trafficking, it may be that the effects of Kv7.5 variants on proteostasis are more muted, especially when Kv7.5 variants are heterozygously expressed. A critical factor, however, is that previous studies in hERG and *SCN5A* have established *z*-scores that are built upon large variant datasets that include benign controls and pathogenic variants with varying severity. In this way, *z*-scores define how variant function deviates from the average function across a large benign population. Assignment of variant pathogenicity as benign, moderately pathogenic, or severely pathogenic is further guided by clinical data with a quantifiable, measurable outcome measure (e.g. the corrected QT interval in the electrocardiogram) to define clinical manifestation. While our development of a *z*-score system establishes a first approach at deriving a similar system, it is currently limited by the paucity of benign variants and the correlation with readily definable clinical severity measures. Though we see variants that have ∼50% functional deficit in current density (e.g. V145G^+/-^, W303C^+/-^), this doesn’t translate to two standard deviations away from normal function (*z*-score-2), because we can only compare with WT control data. The limitation of our current *z*-score implementation is perhaps exemplified by the WT full-block, which causes 87% reduction in current density yet does not come close to reaching a *z*-score value of-2. Without benign variants, our *z*-score range is limited by the inherent variability in our WT dataset. This is more pronounced in the measurement of current density, which shows greater cell-to-cell variability than that of biophysical properties. Further studies investigating a broader set of Kv7.5 variants, with both benign and pathogenic effects, will aid the development of a *z*-score system with improved sensitivity to detect dysfunction. A refined *z*-score system combined with clear clinical metrics of pathogenicity will advance physiological insight and better guide clinical interpretation of functional screening data.

### Potential for targeted pharmacogenetic rescue of function

There is considerable interest in the discovery of novel approaches for targeted correction of Kv7 variant channel function. Our findings indicate that the gain-of-function Kv7.5 variants, G347S^+/-^, L341I^+/-^, and P369R^+/-^ exhibit greater sensitivity to block by the Kv7-spceific blocker, XE991. The greater block in these variants with increased open probability is consistent with the open channel block mechanism described for XE991 (30,31). It is interesting that block was also enhanced in the V145G variant, which exhibits loss of function. This may indicate that the mutation, located in the S1-S2 linker of the channel, stabilizes XE991 at its binding site and/or allosterically modifies the binding site to increase binding affinity. While our findings may indicate a potential pharmacogenetic strategy for the correction of enhanced function, it is challenging to envision selective dose-dependent targeting of gain-on-function variant channel activity in the clinical setting. Moreover, the blocking action of XE991 is not specific for Kv7.5/Kv7.3 channels as it also inhibits Kv7.2/Kv7.3 channels that are a major contributor to neuronal M-current repolarization along with Kv7.5/Kv7.3. Thus, the continued drive to develop Kv7.5 channel-specific compounds likely offers more potential for targeted treatment opportunities.

### Limitations

Our approach to transiently transfect Kv7.3 into stably expressing Kv7.5 cell lines facilitated variant functional assessment by increasing the probability that heteromultimers were formed from co-assembly of Kv7.5 and Kv7.3. However, given that heterotetramers form with variable stoichiometry that is dependent on expression, cells may express channel tetramers with a lower ratio of Kv7.5:Kv7.3 than 2:2. Thus, while our use of a >200 pA peak current cut-off sensitively reports reduced current density in V145G^+/-^ and W303C^+/-^, which is indicative of haploinsufficiency, our data may under-report the effects of these loss-of-function variants on biophysical characteristics. Future studies using cell lines stably expressing both Kv7.5 and Kv7.3 would reveal the role of modified co-assembly and subunit stoichiometry on the functional manifestation of Kv7.5 variants. It would also be interesting to explore the impact of KCNE subunit expression, which are known to modulate Kv7 channel expression and gating in a variable manner depending on stoichiometry, and were not included in the present study.

## Acknowledgements

This study was supported by funding from the Natural Sciences and Engineering Research Council of Canada (grant no. RG-PIN-2026-04365 held by TWC). We thank Dr. Harley Kurata for the kind gift of the Kv7.3/GFP construct and Dr. Alvin Shrier for materials and supplies. We thank Drs. Tim Strassmaier, Chai-Ann Ng, and Jamie Vandenberg for helpful discussions and input on experimental design and analysis.

## Author contributions

DVH – conceptualization, design, data acquisition, data analysis and interpretation of data, drafting of manuscript; SV - conceptualization, design, data acquisition, data analysis and interpretation of data, drafting of manuscript; NG – design, data acquisition, interpretation of data; JL – cell culture; MS – molecular cloning; BK – MaxCyte transient transfection; RKG - MaxCyte transient transfection; DP – design, critical review of manuscript; MP - critical review of manuscript; TWC – project idea, conceptualization, design, supervision, drafting manuscript, data analysis and interpretation.

